# Limits to the inference of gene regulation from bulk tissue expression data

**DOI:** 10.1101/2024.10.24.619521

**Authors:** Ching Pan Chu, Alexander Morin, Paul Pavlidis

## Abstract

**Motivation:** Thousands of studies have used co-expression analysis of bulk tissue samples to probe gene regulation. However, the extent that intracellular regulatory signals are present in these data is unclear. Specifically, we lack clarity of the factors that promote or impede the propagation of intracellular regulatory signals from the single cell level to the bulk tissue level. To bring these issues into focus, we developed a novel computational simulator, grounded in real data, to explore the theoretical relationship between events in single cells and bulk tissue expression profiles, and clarify the conditions required for the propagation of intracellular regulatory signals in complex tissues such as the brain.

**Results:** Our simulator first generates single cell expression profiles and subsequently samples and aggregates these single cells to produce bulk tissue expression profiles. Using this framework, we found that there are very specific and unlikely conditions under which intracellular dynamic regulatory signals can be propagated to the bulk tissue level. For the most part, such regulatory relationships, however strong at the single cell level, are unlikely to be detectable. Our results provide a quantitative explanation for why regulatory network inference from co-expression has proved challenging - even with the assistance of other data modalities - and gives the scientific community a set of tools to further explore these issues in both single-cell and bulk tissue data.

**Availability and implementation:** All relevant data are within the manuscript and supplementary files. The code for all data analyses and generation of figures are available on GitHub (https://github.com/PavlidisLab/coex-simulation). A copy of the data has been deposited in Borealis, the Canadian Dataverse Repository (https://borealisdata.ca/dataset.xhtml?persistentId=doi:10.5683/SP3/2CWXY6).

## Introduction

Co-expression analysis performed using bulk tissue samples has been a primary method for inferring the regulatory circuity that govern expression of genes (Examples: Hartl *et al*., 2021; Pearl *et al*., 2019; Ament *et al*., 2018; Willsey *et al*., 2013; Oldham *et al*., 2008). However, the proposition that such co-expression signals reflect gene regulatory interactions remains largely unvalidated (Garcia-Alonso *et al*., 2019; Djordjevic *et al*., 2014; Marbach *et al*., 2012). Here, we define dynamic regulation to be the temporal change of gene expression within single cells. As such, the ideal co-expression experiment should measure co-variability of gene expressions within single cells across time. While this can be done (Dunlop *et al*., 2008; Chen *et al*., 2022), standard transcriptome-wide methods generally only allow measuring snapshots at the time of sample collection. Co-expression computed across single cells using single cell RNA-seq is thus the most granular form of co-expression in practice. If cells of a given type capture the range of within-cell variability, co-expression across cells would be a proxy for co-expression within cells. However, the majority of published co-expression studies compute co-expression across bulk tissue samples that contain many thousands of cells of different cell types. The relationship between single-cell and bulk tissue co-expression is important to understand if the goal is to infer dynamic regulatory relationships among genes.

In this paper, we propose a framework for considering these questions both conceptually and quantitatively. Conceptually, for the same set of samples, there are at least three distinct levels of resolution at which co-expression could be measured: cross-cell co-expression (xCell), cross-subject co-expression (xSubject) and cross-bulk sample (xBulk) co-expression (Fig. 1). First, bulk tissue samples are generally obtained from different subjects, which introduces subject-level variability that can arise from genetic or environmental differences, within the same cell type. Analyzing variability across subjects (xSubject) is now a standard practice in the field of single cell RNA-seq analysis (Crowell *et al*., 2020).

**Figure 1.**
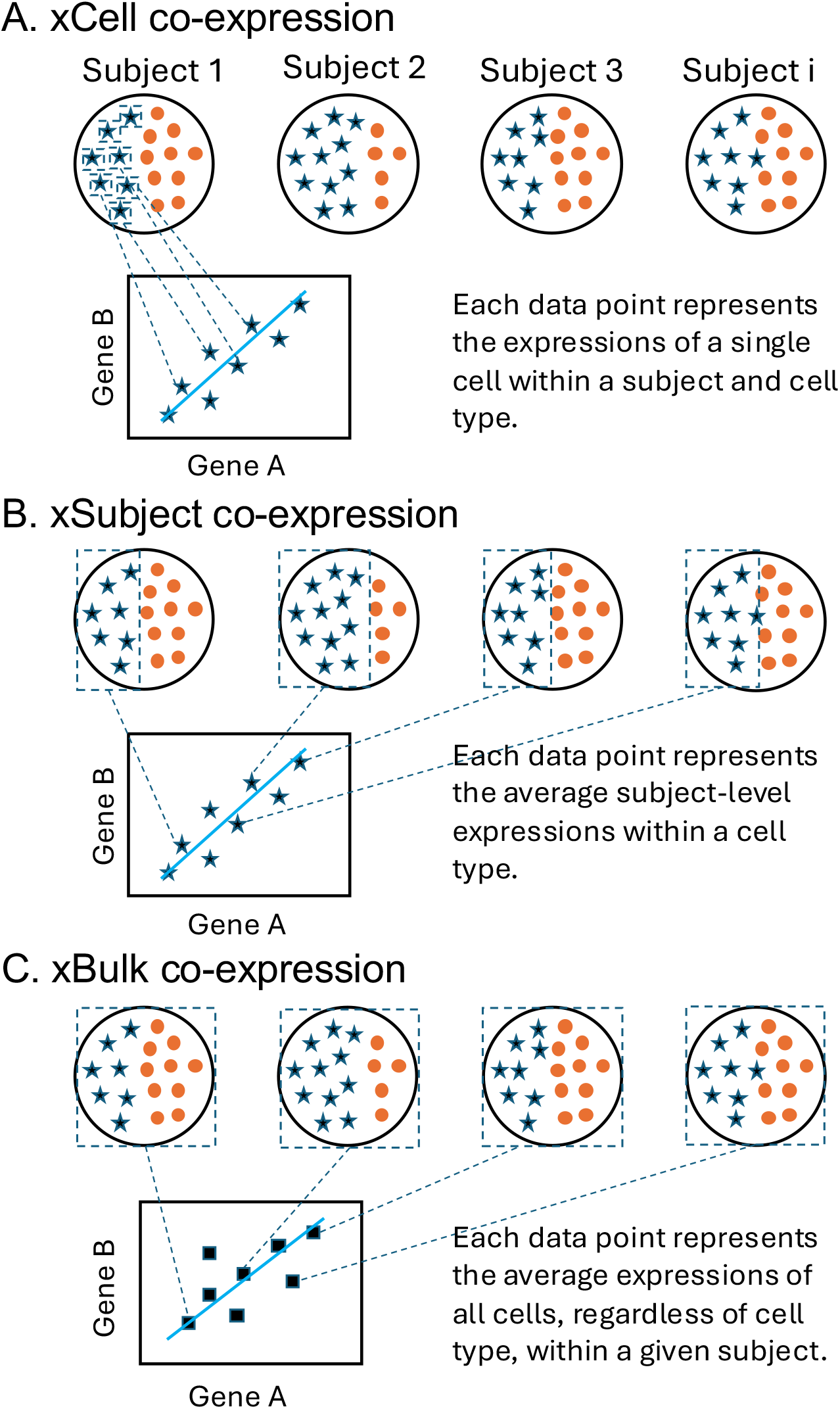
Three levels of co-expression analysis for the same set of biological samples. Schematic depicting the relationship between three levels of resolution where gene co-expression can be computed: xCell, xSubject, and xBulk. (A) At the xCell level, co-expression relationships are computed across single cells of a cell type, within a subject. (B) At the xSubject level, the cell level expression within each subject is first pooled per cell type (i.e. “pseudobulked”), and then co-expressions are computed across the subjects. At the xSubject level, expression patterns are still cell type specific. (C) At the xBulk level, cells of different cell types within each subject are averaged. Co-expressions are then computed across bulk samples that contain different cell types. Only two cell types (stars and circles) are shown in this schematic. The actual number of cell types depends on the tissue in question. In this study, we included six major cell types in the brain (See Methods).

Importantly, co-expression analysis at the xSubject level will expose different types of regulatory signals that may not necessarily align with signals of xCell co-expressions within the same cell type (Supplementary Fig. S1A). To illustrate this, consider genes with sex specific expression. When computed across samples of both sexes, these genes would be strongly positively or negatively co-expressed. However, this xSubject co-expression has no bearing on the genes’ xCell co-expression within the subjects of either sex. Second, bulk samples almost always contain multiple cell types, which may result in the dilution of cell type specific signals (Supplementary Fig. S1B). Third, cellular composition variability (CCV) across bulk samples has been shown to both induce new and mask existing co-expression signals (Farahbod and Pavlidis, 2020) (Supplementary Fig. S1C).

To put these intuitions on a quantitative footing, we use simulations. The use of simulated data allows generation of bulk tissue samples with fully resolved cellular makeup. We developed a novel method for generating single cell expression profiles under a simple model of regulation where co-regulated genes exhibit xCell or xSubject co-expression at the specified level of correlation (Pearson’s r). Subsequently, the simulator samples and aggregates the single cells to produce bulk tissue expressions. Importantly, our approach models single cell expression patterns, and avoids technical sources of variability captured in single-cell genomics experiments, in contrast to the focus of other simulation approaches (Cao *et al*., 2021; Crowell *et al*., 2023). Using this framework, we systematically study the propagation of co-expression signals from xCell, to xSubject, and xBulk levels. Finally, the software that we developed gives the scientific community a set of tools to further explore these issues.

## Materials and Methods

### Reference datasets

We used two reference datasets as training data to estimate parameters for the marginal distributions of gene expression. For the xCell distributions, we used the Allen Institute single-nucleus dataset that was generated using the human middle temporal gyrus (Hodge *et al*., 2019). We selected this Smart-seq dataset for its exceptional sequencing depth. For the xSubject distributions, we used the lymphoblastoid cell lines (LCL) dataset generated as part of the 1,000 genomes project (Lappalainen *et al*., 2013). This dataset enables estimation of xSubject variability within a cell type (See Supplementary Note S1). We mapped the cell type annotations in the Allen Institute data to six broad cell types in the brain: excitatory and inhibitory neurons, astrocytes, microglia, oligodendrocytes, and oligodendrocyte progenitors (OPCs). We filtered the Allen Institute dataset for the cell type and subject with the greatest number of cells (Excitatory neurons: subject H200.1030 with 4,683 cells). For the LCL dataset, we filtered out subjects with duplicated samples, retaining 305 subjects. Finally, we normalized the expression values in both datasets into count-per-million (CPM), which was the intended unit for simulation.

### Simulating expression of genes with co-regulatory relationships

We modeled the marginal distribution across cells and subjects of a given gene’s expression using a mixture of two gamma distributions (Fig. 2A and B; See Supplementary Note S1 for details). We chose the continuous distribution for the convenience and level of control for artificially imposing co-expression structures. In addition, we also avoid dropouts that may be due to technical source of variability. Once constructed, the generative models can be used to produce expressions for any desired number of cells. Briefly, for each gene in the Allen Institute or LCL datasets, we computed the mean and variance values (Fig. 2C). Then, we used a regression model to enable the simulation of xCell or xSubject variance given any mean expression as input. In turn, we used the mean and variance values to parameterize the gamma models. To capture the range of real expression values, we computed the mean expression of each gene in each of the six cell types in the Allen Institute dataset. In total, we included up to 3,000 genes, with 2,880 random genes and 120 marker genes (20 markers for each cell type, defined as genes most specifically expressed in the corresponding cell type). We prefixed the HGNC gene symbols with “sim-” to indicate a gene whose expression profile was simulated. We used the Gaussian copula framework to impose co-expression among genes. The copula offers a way to model marginal distributions of each variable independently from their joint distributions. This method has been used in at least two previous studies for modeling co-expressions (Sun *et al*., 2021; Tian *et al*., 2021).

**Figure 2.**
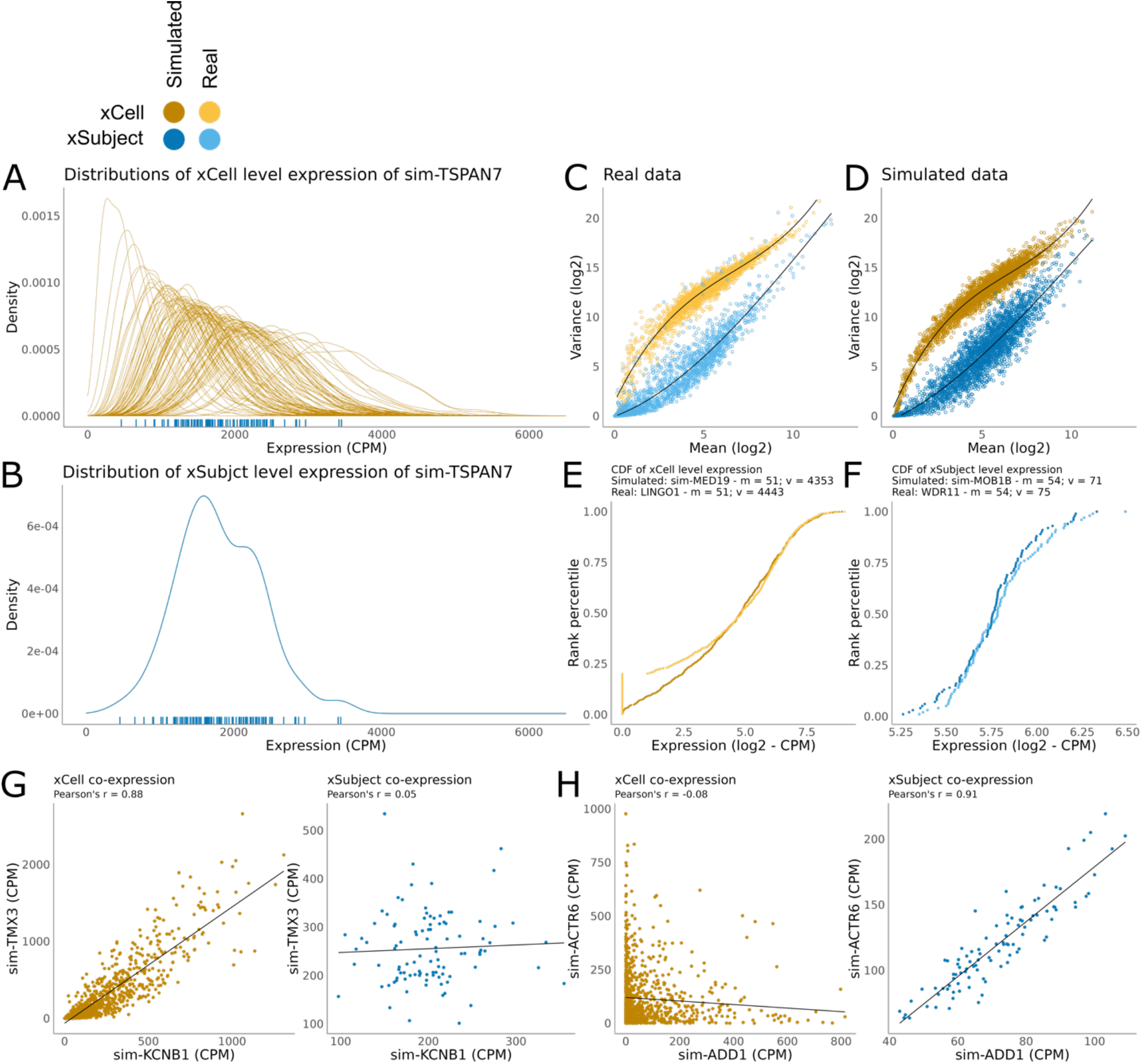
The simulator produces realistic single cell expression patterns with xCell or xSubject co-expression patterns. The color legend for this figure is displayed on the top left corner. (A) Density plots (one per subject) showing the simulated xCell expression distribution of sim-TSPAN7. The mean expression of each xCell distribution is plotted as individual data points on the bottom. (B) The density plot of subject level mean expression of sim-TSPAN7, corresponding to (A, bottom rug plot). (C) Scatterplots showing mean-variance relationship of gene expressions in Allen Institute dataset (excitatory neurons, subject: H200.1030) (xCell, light yellow) and LCL data (xSubject, light blue). Each data point is a gene. (D) Scatterplots showing the mean-variance relationship in the simulated data at the xCell (dark yellow) and xSubject (dark blue). The mean-variance relationship in the real data is recapitulated in the simulated data. (E) The within subject xCell cumulative distribution of sim-MED19 compared to LINGO1 in the Allen Institute dataset. The two genes have comparable mean and variance, but the simulated gene has substantially fewer zeros, as intended. (F) The xSubject level cumulative distribution of sim-MOB1B compared to WDR11 in the LCL dataset. (G) An example of a simulated xCell co-expression (left), which is absent at the xSubject level (right). (H) An example of a simulated xSubject co-expression (right), which is absent at the xCell level (left).

As an initial test of the simulator, we generated an expression dataset containing 1,000 random genes with varying mean and variances in 100,000 cells across 100 subjects for a single cell type: excitatory neurons. Figs. 2C and D show that the mean-variance relationships of the training data were recapitulated at both the xCell and xSubject levels. The marginal distributions of real and simulated data were highly similar for genes with comparable mean and variance values (Figs. 2E and F). Importantly, the simulated cell level expressions no longer contained technical dropouts of lowly expressed genes (Fig. 2E). Finally, Figs. 2G and H show that the simulator was able to impose co-expression relationships separately at the xCell and xSubject levels.

Simulation of subject and bulk level expression profiles are described in detail in Supplementary Note S1. Briefly, each instance of simulation began with the generation of cell level expression profiles for a specified number of cells per cell type, based on realistic estimates (Fig. 3B), across a specified number of subjects. Given the simulated single cell expressions, we computed the subject and bulk level expression profile for each gene by aggregation. Specifically, we derived the subject level expression profiles by averaging the cells for each subject. We modeled bulk level expression of a given gene in a given sample as the weighted sum of the subject level expression of the same gene where the weights were the cell type proportions (Fig. 5A).

**Figure 3.**
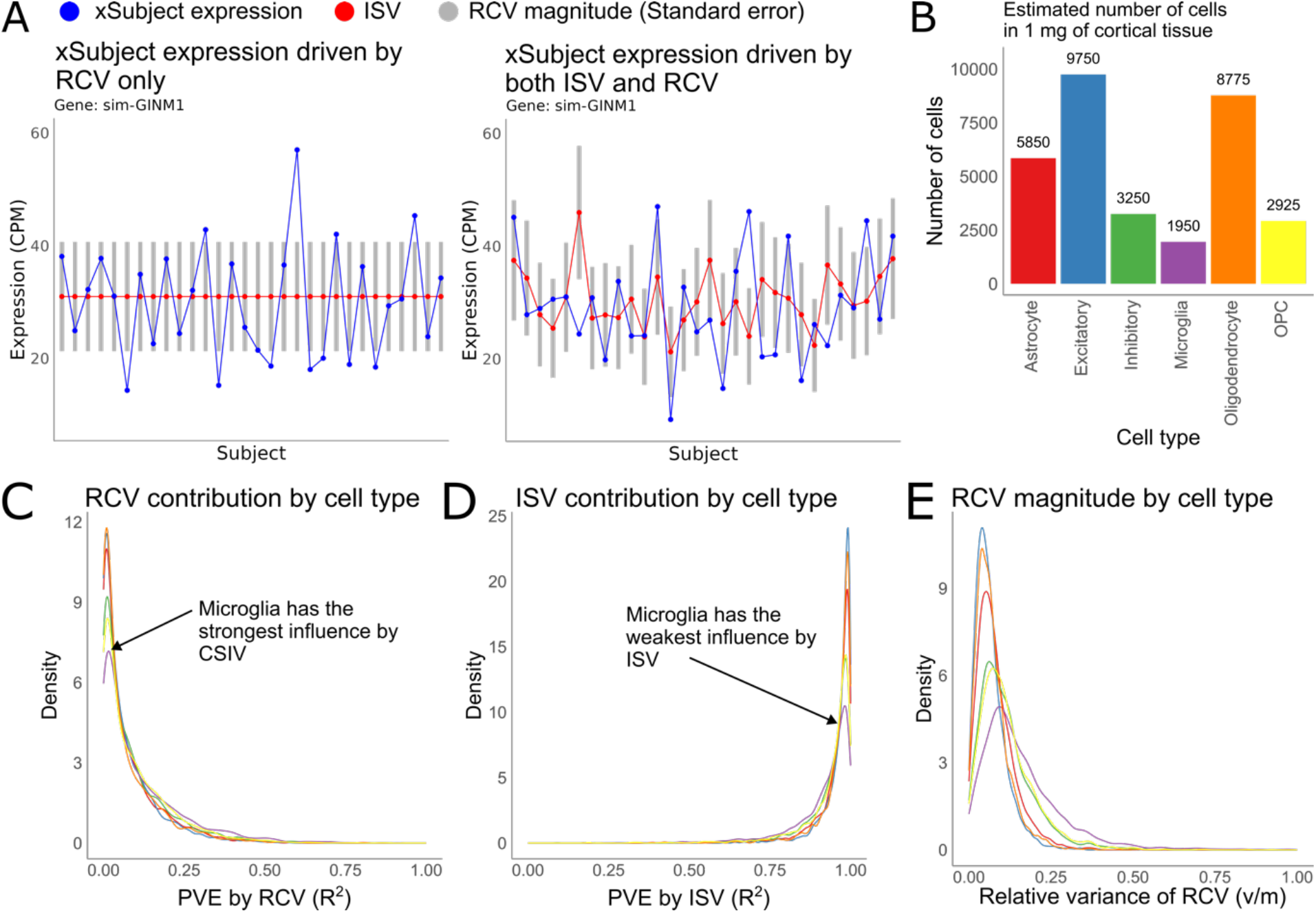
Random cell sampling variability (RCV) is a weak source of xSubject variability compared to intrinsic xSubject-variability (ISV). (A) The RCV expression vector of sim-GINM1, in absence of ISV, is plotted in blue, against the backdrop of the corresponding standard error (gray bars) at 100 cells per sample (left). Note that the true subject level expression, *ISV* of sim-GINM1, is constant across all 30 subjects (red points). The effect of adding ISV is plotted in the panel on the right. (B) Estimated number of cells of different cell types in a 1mg bulk cortical tissue sample. (C) Distribution of proportion of variance explained by RCV across genes, by cell type (30 simulated subjects, 3,000 simulated genes). Each line represents one cell type using the colors as in C. (D) Distribution of proportion of variance explained by ISV across genes, by cell type as in D. (E) Distribution of the magnitude of RCV relative the mean expression for each gene.

## Results

### Cross-cell level co-expression is unlikely to directly propagate to the cross-subject level

Before delving into the role of cell type heterogeneity on co-expression analysis at the bulk tissue level, we first ask a more fundamental question. Within a cell type, can co-expression patterns at the xCell level directly propagate to the xSubject level? This scenario is analogous to sequencing samples that contain purified cell types from different subjects. Since there is a finite number of cells sampled for each subject, the subject level expression is, in effect, a sample estimate of the true mean expression of the given subject. In other words, repeated samples of cells from the same subject could give varying estimates (Figs. 3A; Supplementary Note S2). This variance will go down as counts of cells are increased. We call this “Random Cell-sampling Variability” (RCV). Since RCV is a direct consequence of xCell level variability, it is guaranteed to recapitulate xCell level co-expression patterns. The question is whether this source of variability is large enough to be visible at the xSubject level. The magnitude of RCV is, by definition, the standard error of the xCell distribution, which is inversely related to the square root of the number of cells per sample. As such, RCV will be enhanced by using smaller amounts of source tissue (numbers of cells). Importantly, it is also expected that there would be “Intrinsic Subject-level Variability” (ISV), driven by inter-individual biological differences. As such, RCV must compete with ISV for signal propagation. In effect, the capacity for RCV to drive variability across subjects depends on its magnitude relative to ISV (Fig. 3A).

To explore the relative contributions of RCV and ISV to the resulting expression profiles we simulated samples of one milligram of brain cortical tissue (Fig. 3B) from multiple subjects (See Supplementary Note S2 for details). For each gene, we computed the proportion of xSubject variance that could be explained by either RCV or ISV. As expected, microglia, with the smallest number of cells per sample, had the highest contribution by RCV (purple line in Fig. 3C). Nonetheless, we found that RCV constituted a negligible source of variability for most cell types compared to ISV (Fig. 3D). Intriguingly, there were a small number of lowly expressed genes where RCV explained most of the output signal (Fig. 3C). Naively, these genes may represent candidates for the direct propagation of xCell co-expression. However, while the RCV for these genes was sufficiently large to overcome ISV, none of them had relative variances of >1 (Fig. 3E). Because technical variability of sequencing has a relative variance of >=1 (Marioni *et al*., 2008), RCV signals are unlikely to survive sequencing. We also note that the small sample size of one milligram likely yields optimistic (high) estimates for the degree of direct propagation of xCell co-expression to the xSubject level, compared to most real experiments, which typically use at least 10-100mg of tissue. Taken together, observable xSubject level co-expression patterns are likely independent from the xCell level co-expression in most real datasets. The signal might be enhanced if very small numbers of cells are sampled per subject, but likely well below what is recommended or technically feasible.

### In heterogeneous tissue, cell type specific co-expression signals are easily diluted out

In the previous section, we showed that co-expression at the xCell level would not readily lead to co-expression at the xSubject level (via RCV). However, it remains possible for xSubject co-expression to coincide with the xCell level co-expressions at the level of ISV. This can happen if the same regulatory mechanisms confer variability both across cells and across subjects. However, it remains questionable whether such co-expression signals would survive dilution upon mixture with other cell types in bulk tissue data. The relationship between cell type specific xSubject expression profiles and the resulting diluted bulk expression patterns is schematized in Fig. 4A and explained in Supplementary Note S1.

**Figure 4.**
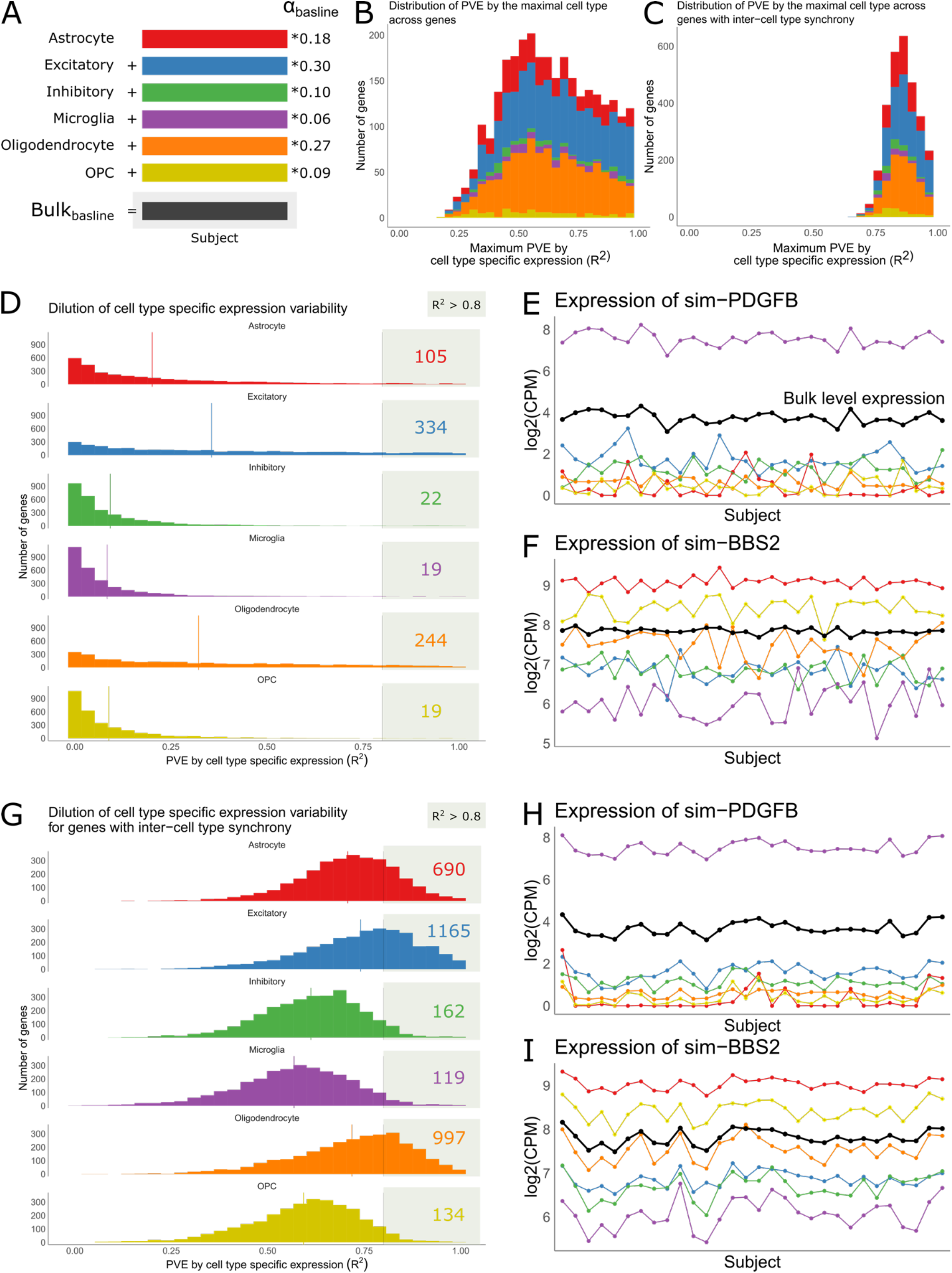
Cell type specific xSubject expression patterns undergo dilution. (A) Schematic showing the relationship between cell type specific xSubject expression vectors and the bulk level expression vector for a given gene. Essentially, bulk level expression is modeled as a weighted sum of the xSubject expressions where the weights, *α*, are cell type proportions (See Supplementary Note S1). To isolate the effects of dilution, cell type proportions are held constant across subjects in this simulation. (B) Distribution of proportion of variance explained (PVE; R^2^) by the maximally explanatory cell type, across genes. For a given gene, a separate *R*^2^ is computed for each cell type specific xSubject vector. The maximum *R*^2^ is plotted here, colored by the corresponding cell type. (C) Same as (B) but for genes with inter-cell type synchrony at Pearson’s r = 0.8. (D) Distribution of PVE, *R*^2^, for each cell type. The lower the PVE, the stronger the effects of dilution on the cell type specific xSubject expression pattern. (E) The xSubject and bulk level expression vectors of the gene sim-PDGFB. In this example, bulk tissue expression of sim-PDGFB (black line) is driven primarily by microglia (purple) due its exceptional cell type specific expression and variability. (F) Same as (E) but for sim-BBS2 whose bulk level expression profile does not resemble any cell type specific xSubject expression vector due to the overall dilution. (G-I) Same as (D-F) but for an alternate simulation where all the genes exhibit inter-cell type synchrony at the xSubject level. This results in the high preservation of any cell type specific xSubject expression, regardless of cell type specificity.

To explore the effects of dilution, we simulated expression data for 3,000 genes and 30 bulk samples. For each gene, we computed the proportion of bulk level variance explained by each cell type specific subject level expression profile. As expected, excitatory neurons, with the highest cell type proportion (30%), also explained the highest amount of variance compared to other cell types (average *R*^2^ = 0.4; Fig. 4D). In contrast, microglia, which only accounts for 6% of the cells in each bulk tissue sample, explain the least amount of variance in the simulated bulk dataset (average *R*^2^ = 0.1; Fig. 4D). By extension, co-expressions in excitatory neurons would also be more preserved than those in other cell types. It is notable, however, that the average preservation of cell type specific variability was generally low. Taken together, these results indicate that most cell type specific expression patterns are expected to be lost by dilution.

The model proposed in Fig. 4A also predicts that propagation of cell type specific variability depends on the cell type specificity of the given gene. If a gene is highly variable only in a one cell type and relatively stable in others, this cell type specific variability would dominate the resulting expression profile even if the cell type in question has a lower proportion than others. Since there is generally a strong mean-variance relationship in expression data (McCarthy *et al*., 2012), we hypothesize that genes with cell type specific expression would be more likely to survive dilution. Consistent with this hypothesis, we observed instances of genes with high degrees of cell type specificity that almost fully retained their cell type specific xSubject expression patterns. This was exemplified by sim-PDGFB in microglia, where the microglia specific xSubject expression explained *R*^2^ = 0.96 of the total variance at the bulk level (Fig. 4E). In total, there were 19 genes in microglia with *R*^2^ values larger than 0.8 (Fig. 4D). By extension, the bulk level co-expression patterns among these genes would directly match those specifically in microglia. There were similar sets of genes in other cell types as well (Fig. 4D). Nonetheless, in this simulation, only 743 (25%) of 3,000 genes were expressed with sufficient cell type specificity required for preservation at *R*^2^ > 0.8 in any cell type, The majority of genes consisted of signals from multiple different cell types, resulting in the general loss of cell type specific expression patterns. This was exemplified by sim-BBS2 (Fig. 4F).

Conceivably, if the expression pattern of a gene in a particular cell type is shared with other cell types, its signal would not only be preserved but reinforced at the bulk tissue level (Fig. 4G to I). We refer to this as “Inter-cell type synchrony”, which can prevent the effects of dilution. For example, circadian rhythms are associated with a daily periodic transcription pattern that is not considered cell-type-specific (Li *et al*., 2013). As a test, we simulated another 3,000 genes that were synchronized among all the different cell types at the xSubject level, at Pearson’s r = 0.8. In this scenario, we observed that the expression patterns of any cell type were largely preserved, for all genes (Fig. 4G). In contrast to genes without inter-cell type synchrony, the preservation of expression patterns included the genes that exhibit no cell type specificity (Fig. 4I).

### Cellular composition variability induces co-expression signals that are independent from intracellular regulation

In the previous section, we assumed that cell type proportions remained constant across bulk samples. In practice, cell type proportions are variable. Previously our lab has shown that this variability is a major driver of co-expression patterns in bulk tissue data (Farahbod and Pavlidis, 2020), a phenomenon referred to as “cellular composition variability (CCV) induced co-expression”. Because CCV-induced co-expression is disconnected from xSubject level regulation within any one cell type, we consider it a source of false positives with respect to the discovery of co-regulation happening within cells. In this section, we simulated 30 bulk samples with varying cell type proportions for 600 genes to further explore the theoretical impact of CCV on expression patterns (See Supplementary Note S1 for details). To produce bulk expression profiles for gene A, denoted *Bulk*_*CCV*_ (*A*), we aggregated the cell type specific xSubject expression vectors as depicted in Fig. 5A. We also computed a version of bulk expression with constant cell type proportions as depicted in Fig. 4A, denoted *Bulk*_*baseline*_ (*A*). For each gene, we used regression models to measure the proportion of variance *R*^2^ in *Bulk*_*CCV*_ that could be explained by the cell type proportions (*α*) versus *Bulk*_*baseline*_(Fig. 5B; Supplementary Note S1).

**Figure 5.**
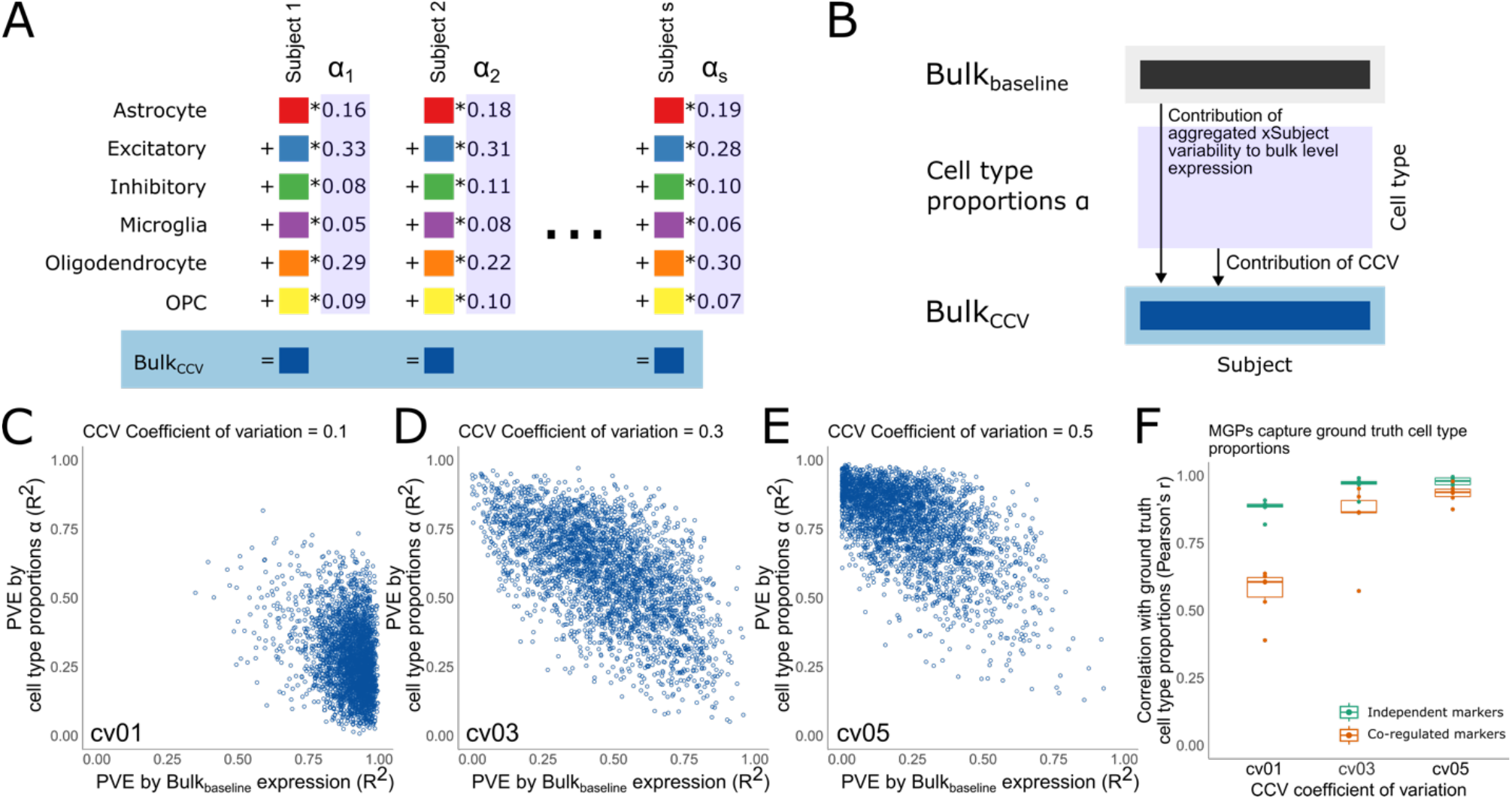
Cellular composition variability (CCV). A) Schematic showing the relationship between xSubject expressions and bulk level expression for a gene in the presence of CCV (Bulk_CCV_). Bulk level expression with CCV is modeled as a weighted sum of the (subject level) expression of the gene in each constituent cell types where the weights are the cell type proportions, *α*. In this simulation, cell type proportion is variable across subjects for each cell type. (B) Schematic showing the dimensions of the cell type proportions matrix and the *Bulk*_*baseline*_ and *Bulk*_(()_expression vectors for a single gene. *Bulk*_2*baseline*_ expression is derived by aggregating the same xSubject expression vectors as in bulk_CCV but without cell type proportion variation. By comparing *Bulk*_*baseline*_ against the cell type proportions and *Bulk*_*baseline*_, we evaluated the contributions of these two sources of variability. (C-E) Scatterplots showing the PVE by *Bulk*_*baseline*_ vs. cell type proportions across 3,000 genes for increasing amounts of CCV. There is an overall negative correlation between the two sources of variability. Importantly, cell type proportions become more important with the magnitude of CCV. (F) Correlations between MGPs and ground truth cell type proportion vectors. MGPs are accurate in absence of inter-marker gene co-expressions (green box plots).

As expected, the explanatory power of cell type proportions increased with the magnitude of CCV variance (Figs. 5C to E). At *cv* = 0.1, cell type proportions could only explain an average of *R*^2^ = 0.28 of the total variance across genes. This increased to *R*^2^ = 0.60 at *cv* = 0.3, and *R*^2^ = 0.77 at *cv* = 0.5. The proportion of variance explained by *Bulk*_*baseline*_ was negatively correlated with that explained by *α* (Figs. 5C to E). CCV increased the average pairwise co-expressions among the genes that were not simulated with co-regulation from Pearson’s r = 0 in *exp*_*baseline*_ to Pearson’s r of 0.06, 0.40, and 0.54 in *Bulk*_*CCV*_ at *cv* of 0.1, 0.3, and 0.5, respectively (Supplementary Fig. S3A). For comparison, we simulated expression for another 600 genes that were co-regulated at Pearson’s r = 0.8. For these genes, the presence of CCV increased the co-expressions slightly relative to *Bulk*_*baseline*_ (Supplementary Fig. S3E). Importantly, the difference between co-regulation and CCV induced co-expression becomes increasingly difficult to distinguish with the increase of the magnitude of CCV.

Next, we explored the feasibility of corrective procedures to remove the effects of CCV. In principle, restoration of cell type specific xSubject expression profiles is impossible. Instead, the best possible performance is the restoration of co-expression patterns that were present in *Bulk*_*baseline*_. Here, we removed CCV as a source of variability by extracting the model residuals, denoted *Bulk*_*residual*_, from the multiple regression models where we previously used cell type proportions (*α*) as covariates to explain variability in *Bulk*_*CCV*_. Reassuringly, following this procedure, the CCV induced co-expressions were removed for all three levels of CCV (Supplementary Fig. S3B). For the co-regulated genes, the original levels of co-expression remained intact (Supplementary Fig. S3F). These observations indicate that CCV correction by regression using known cell type proportions can remove CCV induced co-expression patterns while preserving true regulatory signals.

The above simulation assumes that cell type proportions are known, but in real data these generally must be estimated using deconvolution methods. We thus moved to testing marker gene profiles (MGPs) in place of the known cell type proportions. For this analysis, we selected 20 marker genes for each of the six cell types that exhibited cell type specific expressions in the Allen Institute dataset. We constructed two versions of marker gene profiles for each cell type. In the first version, we constructed the marker gene profiles, denoted *MGP*_*indep*_, using markers that were independent from each other in *Bulk*_*baseline*_. In the second version of MGPs, denoted *MGP*_*coreg*_, we imposed co-regulatory relationships among the marker genes at Pearson’s r = 0.8. Both *MGP*_*indep*_ and *MGP*_*coreg*_ had the same dimensions as cell type proportions *α*, with K rows (cell types) and S columns (subjects) where the *k*^*th*^ row contained the MGP for cell type *k* (Fig. 5B). In both cases, we summarized expressions of the marker genes using principal components analysis and designated the PC1 scores as the MGPs (Mancarci *et al*., 2017).

As an initial test of the method’s capacity to approximate cell type proportions, we computed the correlations between the MGPs and the corresponding cell type proportions *α*_4_ for each cell type *k*. we found that the MGPs in (*MGP*_*indep*_) were highly correlated with the ground truth cell type proportions across all levels of CCV variance (Fig. 5F). Importantly, correction using *MGP*_*indep*_ performed similarly to using the ground truth cell type proportion vectors (Supplementary Figs. S3C and G). However, in *MGP*_*coreg*_, the MGPs had a notable reduction their correlations with the ground truth cell type proportions (Fig. 5F). Correction using *MGP*_*coreg*_ also failed to effectively remove CCV induced co-expressions (Supplementary Fig. S3D). Instead, it removed the co-regulatory signals of genes that shared the same expression patterns as the marker genes (Supplementary Fig. S3H). In conclusion, MGP based CCV correction can be helpful under the condition that marker gene expressions are not co-regulated with each other or with other genes whose expression profiles require correction.

### The fate of co-expression signals from single cell to bulk tissue

So far, we have demonstrated that xCell co-regulatory signals must coincide with xSubject regulation, survive dilution, and distortion by CCV in order to be preserved at the xBulk level. In this final section, we explored five scenarios with varying outcomes of signal preservation. For each scenario, we simulated an expression dataset containing 150,000 cells across 30 subjects (5,000 cells per subject). We computed pairwise co-expressions for all possible pairs of genes at the five different levels of analysis (Fig. 6).

**Figure 6.**
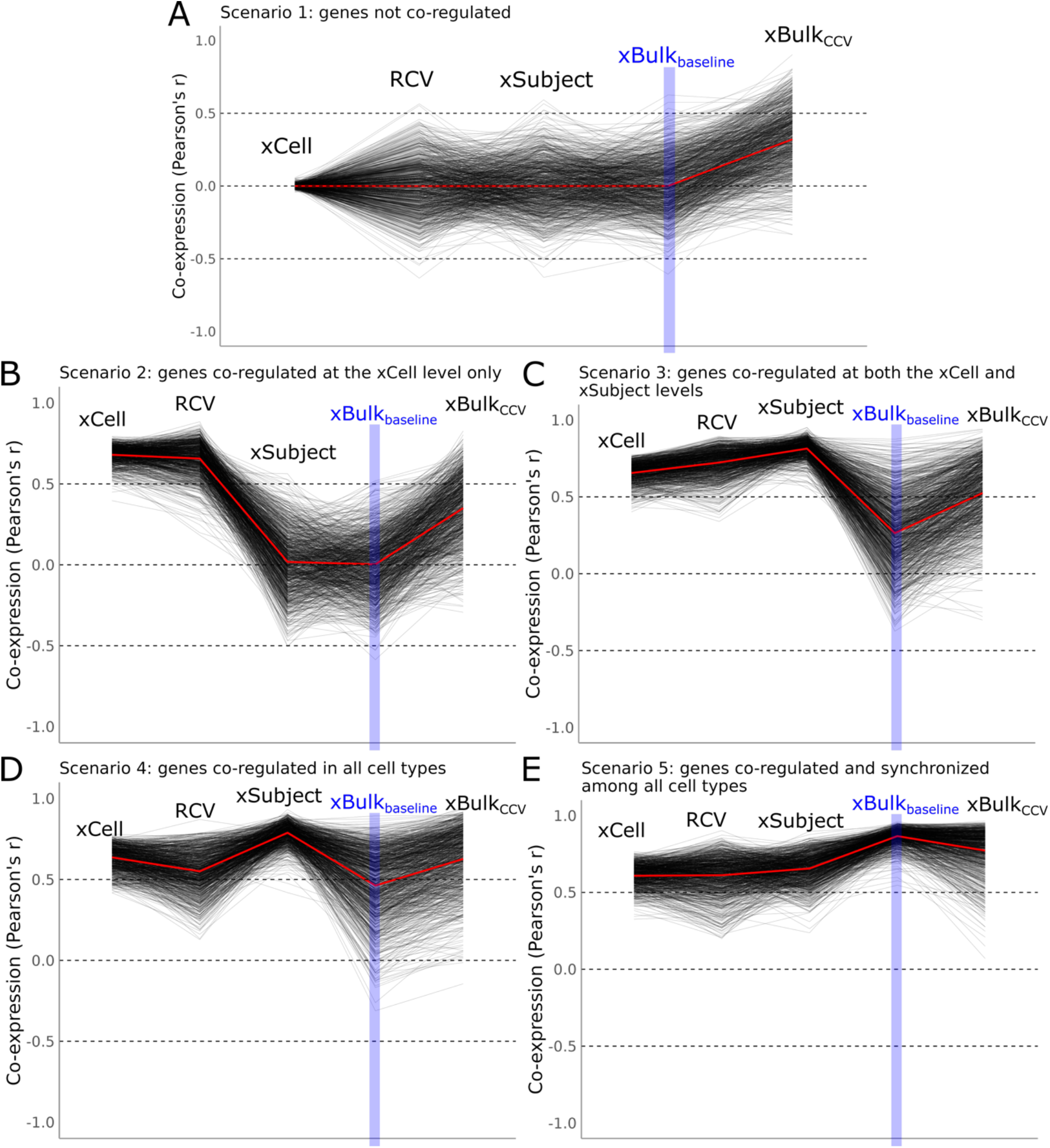
The fate of co-expression signals under different simulation scenarios. The trend charts show the change in pairwise co-expression measures at different levels of cellular resolution. Each line represents the correlation of a pair of genes. The x-axis shows six incremental steps in decreasing levels of resolution, from left to right: 1: xCell is co-expression computed across single cells for excitatory neurons, 2. RCV is co-expression computed across samples in absence of ISV for excitatory neurons, 3. xSubject is co-expression computed across samples with ISV for excitatory neurons, 4. xBulk_baseline_ is co-expression computed across bulk samples containing six cell types of constant proportions, in absence of inter-subject CCV. This level of analysis is highlighted because it contains signals propagated directly from the xSubject level, 5. xBulk_CCV_ is co-expression computed across bulk samples containing variable cell type proportions. The y-axis shows level of co-expression. For the first three levels from left, we plotted the excitatory neuron specific co-expressions. (A) Scenario 1: non-existent co-expression. (B) Scenario 2: co-expression exists at the xCell level only. (C) Scenario 3: co-expression imposed at both the xCell and xSubject levels. (D). Scenario 4: Co-expressions exist independently within all cell types. (E) Scenario 5: co-expression exist within all cell types and all genes were also synchronized among different cell types across subjects.

In Scenario 1 (Fig. 6A), we simulated expression data for 600 genes that do not have true xCell or xSubject co-regulatory relationships, since sampling as well as CCV effects can produce false positives. If false positive co-expressions exceed the strength of co-regulatory signals, they could not be easily distinguished. With a large sample size of 5,000 cells, xCell co-expression of the genes in this scenario centered tightly around Pearson’s r = 0 (Fig. 6A), as expected. Although RCV across the 30 subjects maintained the mean co-expression at Pearson’s r = 0, there was a larger spread of co-expression values due to the smaller sample size. The absence of co-expression signals was recapitulated again at the xSubject level. False positives that arise from sampling could be alleviated by using larger sample sizes. However, the effects of CCV would still induce numerous false positives. This was evident in Fig. 6A, where the distribution of co-expressions underwent a clear shift upwards to a mean Pearson’s r of 0.40 at CCV coefficient of variation *cv* = 0.3. The highest CCV-induced co-expression had a Pearson’s r = 0.92, emphasizing the difficulty of distinguishing such artifacts from true signals of co-regulation.

In Scenario 2 (Fig. 6B), we explored a set of genes with strong (Person’s r = 0.8) xCell co-regulation in excitatory neurons. Importantly, we did not impose co-regulation among these genes at the xSubject level. As expected, RCV directly preserved signals of xCell co-expression. However, since RCV was largely overridden by ISV at the xSubject level, we observed a complete elimination of the xCell co-regulatory signals at the xSubject level (Fig. 6B). Past the xSubject level, there was effectively no difference between the genes in Scenario 2 and Scenario 1. This simulation drives home the point that co-regulation at the xCell level alone would be invisible at the xSubject or xBulk levels.

In Scenario 3 (Fig. 6C), we modeled a set of genes with strong co-expression at both xCell and xSubject levels specifically in excitatory neurons. These co-expression signals were largely lost upon mixture with other cell types. The average co-expression went from Pearson’s r = 0.82 to 0.27 upon dilution. There was a subset of genes whose cell type co-expression patterns were highly preserved (Fig. 6C). As depicted in Section 3.3.3, these signals were mediated by their cell type specific expression. Taken together, these findings indicate that cell type specific co-expressions are in general unlikely to be preserved unless they coincide with cell type specific expression.

It is natural to consider whether co-expression shared by many cell types would be more resistant to the effect of dilution. In Scenario 4 (Fig. 6D), we modeled 600 genes that were highly co-regulated at both the xCell and xSubject levels in all six cell types. While these co-expressions were preserved to a larger extent than the cell type specific co-expressions in Scenario 3, they were still not completely immune to the effect of dilution. On average, co-expression signals of these genes decreased from Pearson’s r = 0.80 to 0.47 after aggregation. The reason for this is that the expression patterns of two co-expressed genes may undergo different effects of dilution due to differences in their cell type expression specificity. In Fig. 4G-I, we showed that inter-cell type synchrony can highly preserve cell type specific xSubject expression patterns of genes upon dilution. Finally, in Scenario 5 (Fig. 6E), we explored the case where genes were not only co-expressed within each cell type, but also synchronized in their expression patterns at the xSubject level at Pearson’s r = 0.80. Compared to other scenarios, this group of genes preserved their co-expression signals across the different stages of cellular resolution more faithfully.

## Discussion

Our study shows that one should not assume bulk tissue co-expression analysis to be informative about dynamic gene regulation. This finding is based on simulations using realistic, or even optimistic, model parameters. Perhaps in hindsight, this finding might be seen as partly intuitive, but we see a main contribution of our study as providing a concrete theoretical framework that has, to our knowledge, been lacking in discussing the information contained in co-expression data. Furthermore, by instantiating this framework in simulation software, we also provide the means for further investigations of different scenarios and parameter settings. Here we summarize and review the implications of our study.

Although there is unlikely any direct propagation of xCell regulatory signals to the xSubject level, it remains possible for xSubject regulation to coincide with xCell regulation. For instance, if the activity of a given transcription factor (TF) is sufficiently variable both across cells within each subject, as well as across subjects, its targets would be detectably co-expressed at both the xCell and xSubject levels. Conversely, if the activity of a given TF is stable across subjects, its targets would be less variable, regardless of their covariance at the xCell level. The key finding that we report here is the theoretical decoupling between xCell and xSubject regulation.

In heterogenous tissues such as the brain, the effects of dilution can be strong. According to our simulations, most the cell type specific co-expression signals would be lost upon aggregation into bulk tissue samples. As such, preservation of co-expression signals requires inter-cell type synchrony at the xSubject level. Inter-cell type synchrony offers an explanation for the preservation of dynamic regulatory signals of circadian rhythm genes at the bulk tissue level (Li *et al*., 2013). The implications of these observations extend naturally to bulk tissue level differential expression analysis. The effects of dilution would make genuine cell type specific expression differences difficult to detect, especially in less abundant cell types. On the other hand, concerted changes in expression among multiple cell types would produce a stronger signal.

Our simulations corroborated previous findings that CCV can be a strong source of variability in xBulk transcriptomes (Oldham *et al*., 2008; Kelley *et al*., 2018; Farahbod and Pavlidis, 2020). This observation supports the use of xBulk co-expression to study cell type specific expression and differential expression patterns (Kelley *et al*., 2018). However, CCV is fundamentally different and cannot be easily disentangled with signals of xSubject regulation. In this study, we also showed that it may be possible to computationally reduce the impact of CCV as a source of variability using MGPs. However, corrective procedures require accurate estimation of cell type proportions, and benchmark studies show that estimation methods have at best an agreement of Spearman’s r ∼ 0.6 with stereology-based estimates (Toker *et al*., 2018; Patrick *et al*., 2020). Importantly, such corrective procedures are ultimately based on the observed expression patterns, which may still carry regulatory information. In the case that marker genes are co-regulated with each other and with other genes, correction can lead to removal of true regulatory signals. Based on this, we propose that marker genes should be chosen to avoid co-expression with other marker genes within the given cell type. And even with CCV correction, the effects of dilution remain a fundamental issue.

Given the challenges of interpretation of bulk tissue transcriptomes, in principle coexpression analysis of single-cell genomics should provide a better window into regulation. But single-cell data currently suffers from what is generally perceived as limitations in data quality compared to bulk tissue, with low sensitivity and measurement accuracy, as well as higher cost and complexity. It is for this reason that we believe bulk tissue studies are still heavily referenced in studies of regulation and even used to validate findings from single-cell data (e.g. Ruzicka *et al*., 2022; Velmeshev *et al*., 2019). Our study shows in quantitative detail that there is no good reason to expect that these data types should generally agree, much less offer similar information on regulation. On a larger scale, it suggests all of the thousands of previous studies claiming to be identifying regulation patterns in bulk tissue should be reinterpreted with our findings in mind.

