## Supplementary Notes for "Limits to the inference of gene regulation from bulk tissue expression data"

### Supplementary Information for
### Limits to the inference of gene regulation from bulk tissue expression data

C. Pan Chu, Alexander Morin, Paul Pavlidis

### Supplementary Note S1: Expression data simulator

#### Data acquisition and preprocessing

We used two reference datasets as training data to estimate parameters for the marginal distributions of gene expression. For estimating the parameters for xCell marginal distributions, we used the Allen Institute single-nucleus dataset that was generated using the human middle temporal gyrus (Hodge *et al.*, 2019). In the main text, we refer to this dataset as the Allen Institute dataset. We obtained this dataset on November 23, 2023, directly from the Allen Brain Atlas website (https://portal.brain-map.org/atlases-and-data/rnaseq#transcriptomics). We selected this Smart-seq dataset for its exceptional sequencing depth. For estimating the parameters for the xSubject marginal distributions, we used the lymphoblastoid cell lines (LCL) dataset generated as part of the 1,000 genomes project (Lappalainen *et al.*, 2013). In the main text, we refer to this dataset as the LCL dataset. This dataset contains transcriptomes of pooled LCL cell lines generated from 462 individuals. The large number of subjects, coupled with the bulk level sequencing protocol from a homogeneous population of cells, enables estimation of xSubject level variability within the same cell type. We obtained the LCL dataset from ArrayExpress (https://www.ebi.ac.uk/biostudies/arrayexpress/studies/E-GEUV-1) on November 23, 2023.

For both datasets, we mapped the genes to Ensembl identifiers using HUGO Gene Nomenclature Committee (HGCN) gene symbols. We obtained the Ensembl gene annotation file from Gemma (Lim *et al.*, 2021). Subsequently, we filtered out genes that were detected (>0 sequencing reads) in less than 5% of all samples in each dataset. As a result, we retained 12,164 genes in the Allen Institute dataset and 30,059 genes in the LCL dataset. In the metadata of the Allen Institute dataset, we mapped the cell type annotations to six broad cell types in the brain: excitatory and inhibitory neurons, astrocytes, microglia, oligodendrocytes, and oligodendrocyte progenitors (OPCs). To estimate within cell type xCell variability, we further filtered the Allen Institute dataset for the cell type and subject with the greatest number of cells (Excitatory neurons: subject H200.1030 with 4,683 cells). For the LCL dataset, we filtered out subjects with duplicated samples, retaining 305 subjects. Finally, we normalized the expression values in both datasets into CPM, which was the intended unit for simulations as well.

#### Estimating model parameters

We approached the simulation by constructing generative models based on the observed distributional properties of reference datasets. We modeled the marginal distribution of a given gene's expression using a mixture of two gamma distributions (Fig. 2A and B). Once constructed, the generative models can be used to produce expressions for any desired number of cells. For clarity, each gene has a probabilistic model that defines its distribution across cells and subjects. Specifically, for gene A, its expression within the cells of a given subject $s$, denoted $Exp\_cel_s(A)$, follows a gamma distribution:

$$Exp\_cel_s(A) \sim gamma(k\_cel_s, \theta\_cel_s)$$

where $k\_cel_s$ and $\theta\_cel_s$ denote the subject $s$ specific xCell level shape and scale parameters. The subject level expression of gene A, denoted $Exp\_sbj(A)$, in turn, follows another gamma distribution:

$$Exp\_sbj(A) = E\left[Exp_{cel_s}(A)\right] \sim gamma(k\_sbj, \theta\_sbj)$$

where $k\_sbj$ and $\theta\_sbj$ denote the xSubject level distribution parameters.

We estimated the xCell level model parameters using the Allen Institute dataset and the xSubject level model parameters using the LCL dataset. First, we used a third order polynomial regression model to enable generating xCell variance based on input mean expression. Specifically, for each gene in the Allen Institute dataset, we computed the mean and variance values (Fig. 2C, yellow). Then, we modeled the level of variance of gene A as a function of its mean expression as follows:

$$v_A = f(\mu_A) = \beta_0 + \beta_1\mu_A + \beta_2\mu_A^2 + \beta_3\mu_A^3 + \epsilon_A,$$

$$\epsilon \sim normal(0, \sigma)$$

$$\text{where } v_A = log_2(Var(A)),$$

$$\text{and } \mu_A = log_2(Mean(A))$$

where $\mu_A$ denotes the mean expression of gene A, $v_A$ denotes the corresponding variance, and the $\beta$'s denote the model coefficients. The error term $\epsilon_A$ was modeled using a normal distribution with the standard deviation $\sigma$ estimated using the model residuals. This regression model enables the simulation of xCell variance given any mean expression as input. We repeated this procedure on the LCL dataset to construct a separate regression model to predict xSubject variance. Once generated, the mean and variance values were then used to estimate the shape $k$ and scale $\theta$ parameters for the corresponding gamma distributions using the method of moments:

$$k = \frac{\mu^2}{v} \text{ and } \theta = \frac{v}{\mu}$$

Taken together, these regression models enabled the generation of two sets of gamma model parameters for each gene, with the same input mean but different simulated variances at the xCell and xSubject levels. Integration of the xCell and xSubject models into a single mixed model was performed in two steps. For each gene A, we simulated the subject level expressions using the xSubject distribution model, producing an expression vector $Exp\_sbj(A)$ of size $S$, where $S$ is the number of subjects. For each subject $s$, we constructed a new xCell model by converting the original xCell level shape parameter $k$ to adopt the mean expressions specified by $Exp\_sbj(A)$, while preserving the scale parameter $\theta$ to maintain the model's relative variance. This procedure enabled the construction of a generative model given any arbitrary mean expression as input. To capture the range of real expression values, we computed the mean expression of each of the six cell types in the Allen Institute dataset. In total, we included 3,000 genes, including 2,880 random genes and 120 marker genes (20 markers for each cell type, defined as genes expressed in only that cell type). We varied the actual number of genes in each instance of simulation due to constraints in computational resources by randomly subsampling. The numbers of genes were chosen to sufficiently capture the range of expression levels in real RNA-seq datasets. For each gene and cell type, we constructed a generative model based on the cell type specific mean expression. In total, we constructed 3,000 genes * 6 cell types = 18,000 gamma mixture models for simulation. For all analyses, we included both non-marker and marker genes, unless specified otherwise.

#### Simulating expression of genes with co-expression relationships

The probabilistic models discussed so far only enable simulations of independently distributed gene expressions, without any control over correlation among genes. We therefore used the Gaussian copula framework to impose co-regulatory relationships among genes, which take the form of pairwise correlations in expression profiles. This method has been used in several previous studies for modeling co-expressions (Sun *et al.*, 2021; Tian *et al.*, 2021). The copula offers a way to model marginal distributions of each variable independently from their joint distributions. To simulate the co-expression between genes A and B while preserving their respective marginal distributions, we began by using the mvtnorm (Genz and Bretz, 2009) package in R to generate two correlated vectors, $Exp(A)$ and $Exp(B)$ whose marginal distributions followed the standard normal distribution. Next, we converted the values within each vector to uniformly distributed p-values by using the cumulative distribution function for the standard normal distribution. This step converted the joint normal distribution of $Exp(A)$ and $Exp(B)$ into a Gaussian copula. As a final step, we converted each expression vector into the target marginal distribution models by using the inverse cumulative distribution functions corresponding to the specified gamma models. As a result, $Exp(A)$ and $Exp(B)$ retained their co-expression relationship while following different marginal distributions. Importantly, we were able to use this strategy to impose co-expression relationships separately at the xCell and xSubject levels (Figs. 2G to H).

#### Simulating cell, subject, and bulk tissue level expression data

Each instance of simulation began with the generation of cell level expression profiles for a specified number of cells. To mimic real bulk tissue data, we estimated the cell numbers for each cell types in one milligram (mg) of cerebral cortical tissue by reviewing previous literature. Note that a typical bulk RNA-seq protocol requires 500 to 1,000 nanograms (ng) of purified total RNA, which may be obtained using one to two milligrams of human brain tissue (Walker *et al.*, 2016), though this is much less tissue than typically used for RNA extraction in practice. Based on the meta-analysis done by (Herculano-Houzel, 2009), there are, on average, 13,000 neurons in one milligram of the human cerebral cortex. About 25% percent of all neurons are inhibitory neurons and the remainder are excitatory neurons (Hendry *et al.*, 1987). The number of glial cells is about 1.5 times (19,500) the number of neurons, split into 5,850 (30%) astrocytes, 1,950 (10%) microglia, 8,775 (45%) oligodendrocytes and 2,925 (15%) oligodendrocyte progenitor cells (von Bartheld *et al.*, 2016).

Given the simulated single cell expressions, we computed the subject level expression profile for each gene by aggregation. Within a cell type, the expression of gene $A$ in subject $s$ can be represented as a vector $Exp\_cel_s(A)$ of size $C$ where $C$ is the number of cells. The subject level expression of A is simply $Mean(Exp\_cel_s(A))$. Repeating this calculation for all S subjects within cell type $k$ produces a subject-level expression vector, $Exp\_sbj_k(A)$, of size S. Likewise, repeating this calculation for all the cell types produces six cell type specific xSubject expression vectors. The xSubject expression vectors were, in turn, further aggregated to produce bulk tissue samples containing mixed cell types. Specifically, for a given subject $s$, the bulk tissue expression $Exp\_bulk_s(A)$ is derived as follows:

$$Exp\_bulk_s(A) = \sum_k \alpha_{k,s} * Exp\_sbj_{k,s}(A)$$

where $k$ denotes cell type, $\alpha_{k,s}$ denotes the proportion of cell type $k$ in subject $s$ and $Exp\_sbj_{k,s}(A)$ denotes the subject level expression of cell type $k$. To explore the effects of dilution, we simulated bulk tissue data with no CCV across subjects where $\alpha_{ks}$ is constant within $k$ (Fig. 4A). To explore the effects of CCV, we varied $\alpha_{k,s}$ across the subjects (Fig. 5A). Based on immunohistochemistry (IHC) imaging data provided by (Patrick *et al.*, 2020), cell type proportions vary in cortex by a coefficient of variation $cv$ between 0.1 and 0.3. We also included a stronger case of $cv = 0.5$ for comparison. To generate varying cell type proportions across samples, we modeled the proportion of each cell type as

$$\alpha_k \sim normal(\alpha\_baseline_k, \alpha\_basline_k * cv)$$

where $\alpha_k$ is the cell type proportion of cell type $k$ and $\alpha\_baseline_k$ denotes the baseline proportion of $k$. Using this model, we were able to generate cell type proportion vectors of any length with baseline proportions as input.

#### Cellular composition variability correction

When relevant, to simulate measuring and correcting for CCV effects, we performed CCV correction by using a multivariate regression model to predict bulk tissue expression as a function of cell type proportions. Specifically, cell type proportions can be captured using six vectors, with each vector $\alpha_k$ containing the proportion of cell type $k$ across S subjects. Then, we modeled bulk tissue expression of gene A, $Exp\_bulk_s(A)$, as

$$Exp\_bulk_s(A) = \beta_0 + \sum_k \beta_k * \alpha_{k,s} + \epsilon_{A,s}$$

where $\alpha_{k,s}$ denotes the proportion of cell type k in subject s, $\beta's$ are the model coefficients, and $\epsilon_{A,s}$ denotes the residual error. This model allowed me to measure the proportion of variance, $R^2$, in the bulk expression of A that can be explained by CCV. Importantly, we can remove the putative effects of CCV by extracting the residual of the regression model, thereby accomplishing CCV correction (Farahbod and Pavlidis, 2020). In addition, we also used the marker gene profiles (MGPs) in place of the cell type proportions to accomplish expression-based correction. We estimated MGPs using the method developed by (Mancarcwe et al. 2017). Briefly, for each cell type, we summarized the expressions of the corresponding marker genes by PCA. Finally, we used the PC1 scores as MGPs. To use the MGPs for CCV correction, we substituted the $\alpha_{k,s}$ values with expression based MGPs in the above model.

### Supplementary Note S2: Computing contributions of RCV versus ISV

Starting with excitatory neurons, we simulated expression data for 3,000 random genes and 30 subjects where each subject has 100 cells. In the first version of the simulation, we initially generated cell-level expression profiles, which we denote $Exp\_RCV\_Cel$, where the true mean expression for each gene was constant across all 30 subjects (Supplementary Figure S2B). In other words, the xCell level distributions were identical among all subjects. Using these simulated cells, we derived an expression matrix $Exp\_RCV$ containing 3,000 rows (genes) and 30 columns (subjects) where each element is the average expression of all the cells contained in a given sample. Under this condition, RCV (Random Cell-sampling Variability, see Main Text) is the only source of variability across samples in $Exp\_RCV$. As a second step, we simulated a set of 30 subject level expression profiles, denoted $Exp\_ISV$ with the same dimensions. Each element in $Expr\_ISV$ is the true subject level mean expression of each gene in each subject.

To simulate the final subject-level expression matrix, denoted $Exp\_sbj$, which integrates both RCV and ISV, we applied a transformation to $Exp\_RCV\_Cel$ using the Gaussian copula framework. Specifically, for each gene $g$ and subject $s$, we first derived a new target xCell expression distribution, $gamma(k\_cel_s, \theta\_cel_s)$ where the mean of this target distribution was set by $Exp\_ISV$ while maintaining the relative variance observed in our reference xCell data (as detailed in Supplementary Note S1). With these new parameters, we "shifted" the $Exp\_RCV\_Cel$ values (which embody the RCV characteristics) to conform to these newly defined subject-specific xCell distributions. This is done by using the copula. More specifically, the original values were converted into a uniform distribution using the cumulative distribution function of original xCell distribution and then mapped to the target distribution by using the corresponding inverse cumulative distribution function (See Supplementary Note S1). This transformation ensures that the deviations from the mean due to cell sampling (RCV) are maintained, effectively integrating RCV and ISV in the final subject-level expression matrix $Exp\_sbj$ (Supplementary Figure S2C).

For each gene, we computed the proportion of variance in $Exp\_sbj$ that could be explained by either $Exp\_RCV$ or $Exp\_ISV$. Specifically, for each gene g, we calculated the squared Pearson correlation coefficient ($R^2$) between the combined $Exp\_Sbj(g)$ vector (across subjects) and the pure $Exp\_RCV(g)$ vector (representing variability solely from random cell sampling) or the pure $Exp\_ISV(g)$ vector (representing true subject-level biological differences). The proportion of variance explained by RCV ($PVE\_RCV$) for gene g was calculated as:

$$PVE\_RCV(g) = PearsonCor(Exp\_sbj(g), Exp\_RCV(g))\text{^}2.$$

Similarly, the proportion of variance explained by ISV ($PVE\_ISV$) for gene g was calculated as:

$$PVE\_ISV(g) = PearsonCor(Exp\_sbj(g), Exp\_ISV(g))\text{^}2.$$

As a check, we confirmed that RCV and ISV represent competing signals whose contributions were negatively correlated. At 100 cells per sample, $Exp\_RCV$ explained $R^2 = 0.44$ of the total variance (average across 3,000 genes). As previously mentioned, the magnitude of RCV decreases with increasing numbers of cells per sample. Consistent with this hypothesis, the average proportion of variance explained by $Exp\_RCV$ went down to $R^2 = 0.14$ when we repeated the above experiment with 1,000 cells per sample (Supplementary Figs. S2D to F). Under this scenario, the variability of most genes was dominated by ISV. Taken together, we have shown that RCV, which is a mediator for the direct propagation of xCell co-expressions to the xSubject level, becomes weaker with higher number of cells per sample as expected. Consequently, having fewer cells per sample may improve the visibility of dynamic regulatory signals at the xBulk level.

### Supplementary Figures

#### Supplementary Figure S1

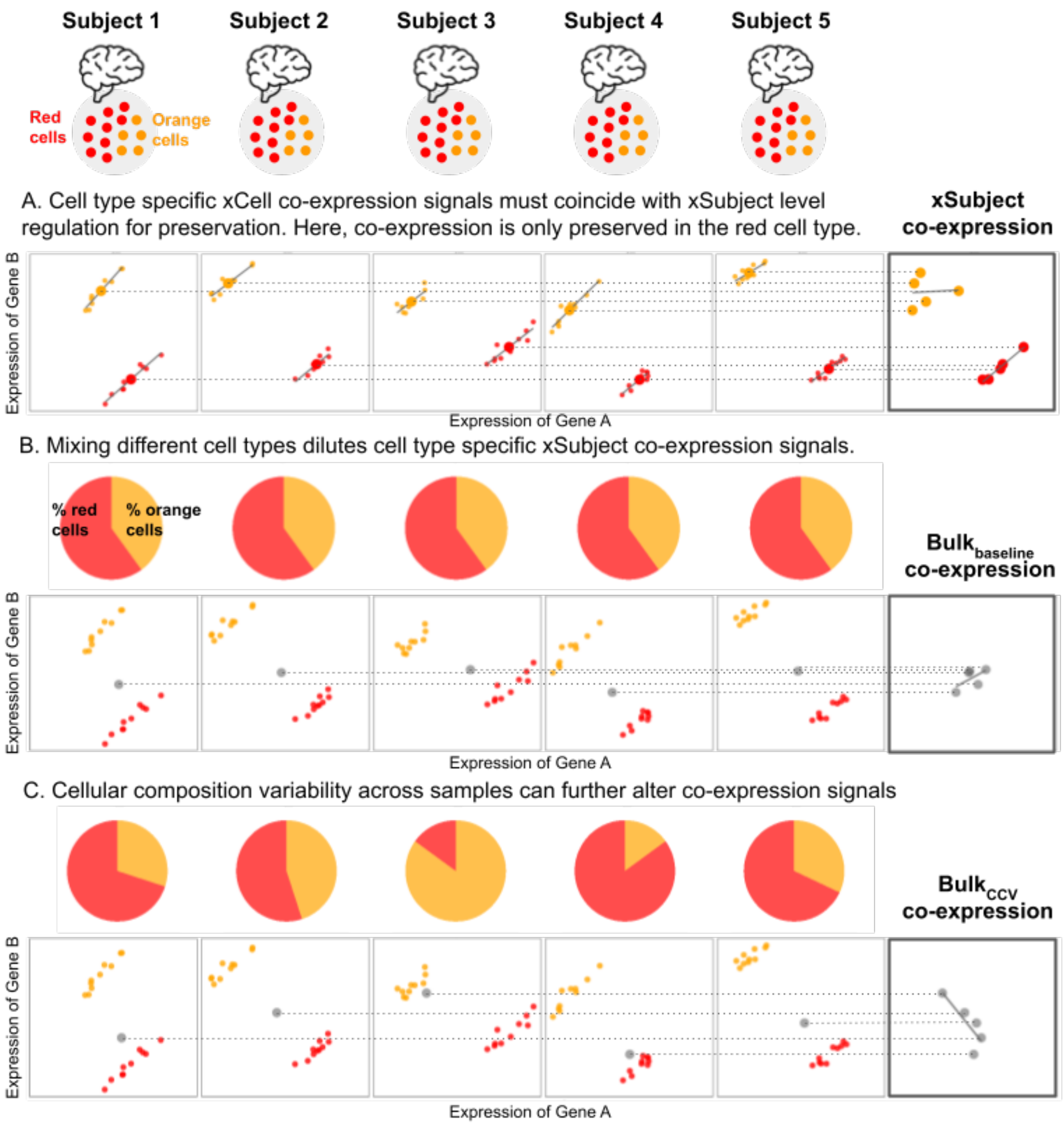


**Supplementary Figure S1.** Concordance of co-expression patterns among different levels of analysis is not guaranteed. Schematic showing the hypothetical scenarios of concordance in the co-expression patterns among the three levels of cellular resolution. (A) Genes A and B are highly co-expressed at the xCell level in both orange and red cell types. However, co-expression is only retained at the xSubject level in red cells but lost in the orange cells. (B) After aggregation of the two cell types at constant cell type proportions, the xSubject level co- expressions in the red cell type are diluted by the absence of co-expression in the orange cell type. (C) Variability in cell type proportions across subjects induces a negative xBulk level co-expression between genes A and B due to their opposite cell type expression specificities. At this level, there is no longer any indication of positive xCell co-expression within either cell type. This figure uses icons created by DinosoftLabs on www.flaticon.com (Flaticon license).

### Supplementary Figure S2

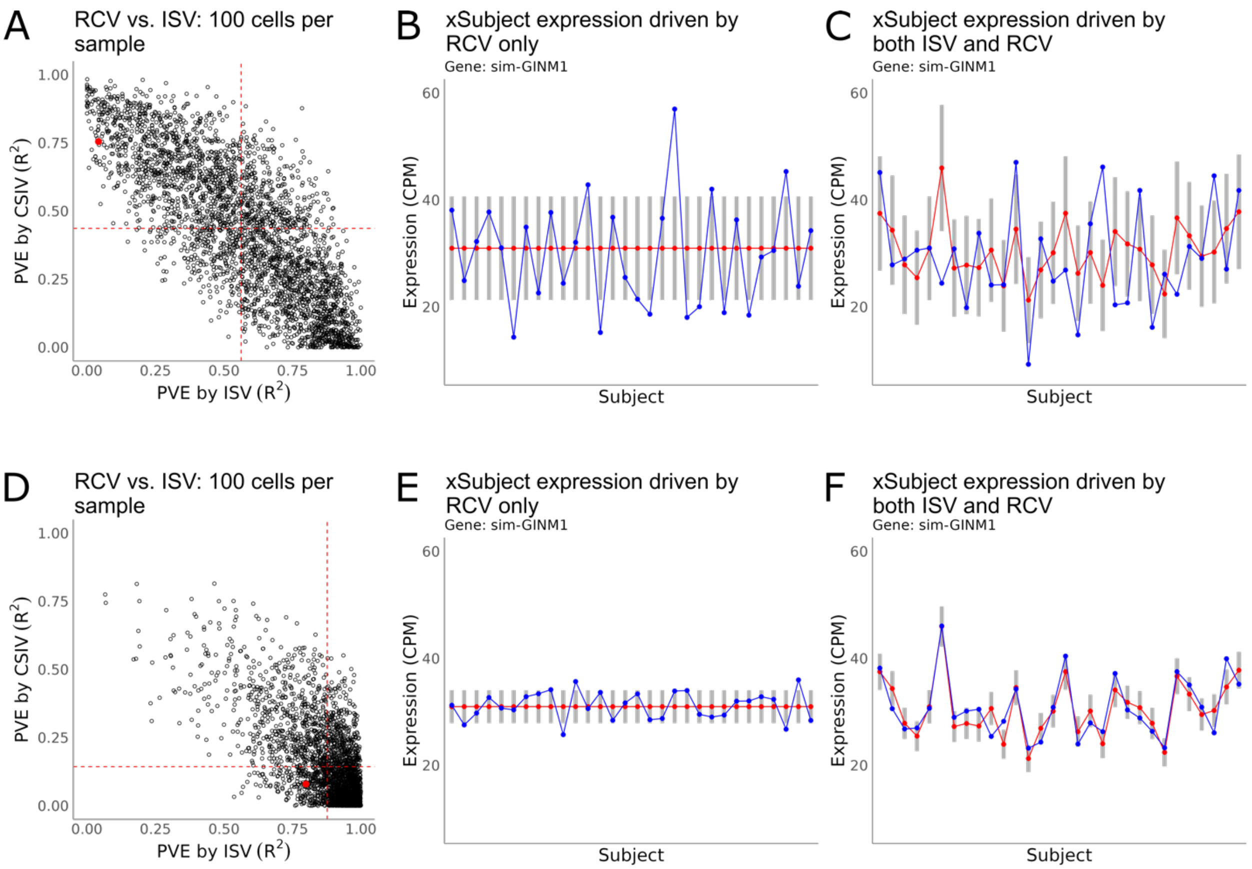


**Supplementary Figure S2.** Random cell-sampling variability (RCV) is a weak source of xSubject variability compared to intrinsic subject variability (ISV). (A) Scatterplot showing the contributions RCV versus ISV to the resulting xSubject level expression patterns at 100 cells per sample. Each data point is a gene. The gene sim-GINM1, highlighted in red, exemplifies a case where xSubject variability is largely driven by RCV. (B) The $Exp\_RCV$(sim-GINM1) vector, in absence of ISV, is plotted in blue, against the backdrop of the corresponding standard error (gray bars) at 100 cells per sample. Note that the true subject level expression, $Exp\_ISV$(sim-GINM1), remains constant across all 30 subjects. (C) The $Exp\_sbj$(sim-GINM1) vector is plotted in blue, which is the result of incorporating $Exp\_RCV$(sim-GINM1) and $Exp\_ISV$(sim-GINM1). Because the standard error is large at 100 cells per sample, RCV remains the dominant signal in this scenario. (D) Same as (A) but for 1,000 cells per sample. (E) Same as (B) but for 1,000 cells per sample. Note that the standard error, and consequently the magnitude of RCV is much smaller compared to (E). (F) Same as (C), but for 1,000 cells per sample. Note that ISV is now a much stronger source of variability.

### Supplementary Figure S3

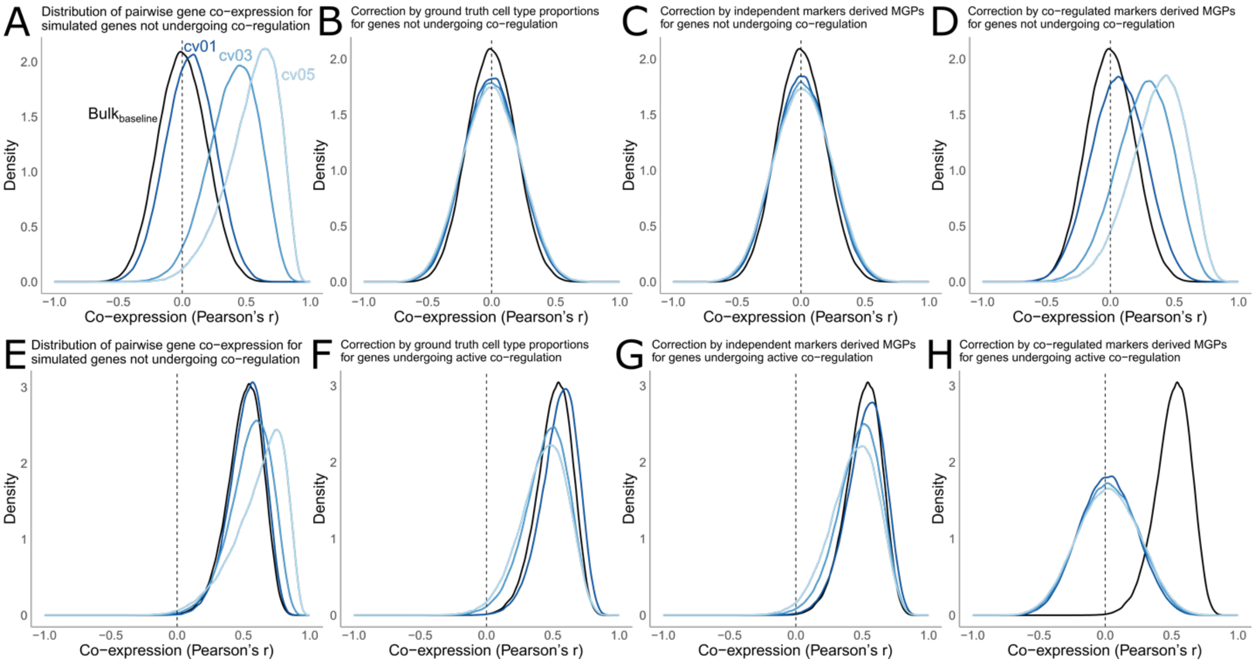


**Supplementary Figure S3.** In-silico correction of cellular composition variability (CCV) in co-expression. (A) Distribution of co-expressions in $Bulk_{baseline}$ and $Bulk_{CCV}$ at the different levels of CCV, in absence of true regulation, prior to any correction. Positive correlations are induced by CCV (B) $Bulk_{CCV}$ expressions were corrected using ground truth cell type proportions, restoring the distribution to baseline. (C) $Bulk_{CCV}$ expressions were also effectively corrected using marker genes with no inter-marker gene co-expressions. (D) $Bulk_{CCV}$ expressions were corrected using marker genes with inter-marker gene co-expressions at Pearson's r = 0.8, resulting in reduced effectiveness of the correction. (E-H) Same as (A-D) but for a group of genes with true co-expressions at Pearson's r = 0.8.
